# Identifying drug-resistant individual cells within tumors by semi-supervised transfer learning from bulk to single-cell transcriptome

**DOI:** 10.1101/2024.08.18.608421

**Authors:** Kaishun Huang, Hui Liu

## Abstract

The presence of pre-existing or acquired drug-resistant cells within the tumor often leads to tumor relapse and metastasis. Single-cell RNA sequencing (scRNA-seq) enables elucidation of the subtle differences in drug responsiveness among distinct cell subpopulations within tumors. A few methods have employed scRNA-seq data to predict the drug response of individual cells to date, but their performance is far from satisfactory. In this study, we propose SSDA4Drug, a semi-supervised few-shot transfer learning method for inferring drug-resistant cancer cells. SSDA4Drug extracts pharmacogenomic features from both bulk and single-cell transcriptomic data using semi-supervised adversarial domain adaptation. This allows us to transfer knowledge of drug sensitivity from bulk-level cell lines to single cells. We conduct extensive performance evaluation experiments across multiple independent scRNA-seq datasets, demonstrating SSDA4Drug’s superior performance over current state-of-the-art methods. Remarkably, with only one or two labeled target-domain samples, SSDA4Drug significantly boosts the predictive performance of single-cell drug responses. Moreover, SSDA4Drug accurately recapitulates the temporally dynamic changes of drug responses during continuous drug exposure of tumor cells, and successfully identifies reversible drug-responsive states in lung cancer cells, which initially acquire resistance through drug exposure but later restore sensitivity during drug holidays. Also, our predicted drug responses consistently align with the developmental patterns of drug sensitivity observed along the evolutionary trajectory of oral squamous cell carcinoma cells. In addition, our derived SHAP values and integrated gradients effectively pinpoint the key genes involved in drug resistance in prostate cancer cells. These findings highlight the exceptional performance of our method in determining single-cell drug responses. This powerful tool holds the potential for identifying drug-resistant tumor cell subpopulations, paving the way for advancements in precision medicine and novel drug development.

## Introduction

Tumor heterogeneity often leads to substantial differences in patient responses to the same drug treatment [1]. Despite remarkable progress in anti-cancer therapy, the presence of pre-existing or acquired drug-resistant cells within the tumor survive and proliferate, ultimately leading to tumor relapse and metastasis [2]. Drug resistance is a leading cause of clinical failure of tumor therapy. Some scholars have developed bioinformatic methods to predict patients’ clinical responses to drugs based on bulk RNA-seq data from cell lines, leveraging large-scale drug sensitivity data such as from GDSC [3, 4], CCLE [5, 6] and CTRP [7]. For example, Chawla et al. introduced the Precily [8] framework that predicts drug response in patients and patient-derived xenografts (PDX) by integrating drug molecular structures and pathway enrichment scores. Meanwhile, He et al. introduced the context-aware deconfounding autoencoder (CODE-AE) [9], which transfers knowledge learned from cell lines to tumor tissues and deconfounds confounding factors involved in drug responses. Additionally, Ma et al. present a few-shot learning for drug response prediction that can translate from high-throughput screens to individual patients [10]. Despite the potential, bulk RNA-seq fails to discern variations in transcriptional programs and regulatory mechanisms within individual cells, thereby obscuring the intratumor heterogeneity. Consequently, previous models perform poorly on out-of-distribution samples, including patients and individual cells.

The advancement of single-cell sequencing technology enables us to explore the complexity and variability among individual cells, offering us the opportunity to gain insights into the intratumor heterogeneity [11]. However, the currently available drug response data at single-cell level covers only a few cancer types and drugs due to cost and technical limitations [12], which poses challenges on the development of computational methods for predicting single-cell drug responses. Several studies have focused on the knowledge transfer of drug responses from cell lines to single cells. The unsupervised domain adaptation technique, including maximum mean discrepancy [13–15], adversarial learning [16–19], or reconstruction learning [20, 21], has been used to align the data distribution between the source and target domains. For example, scDeal [22] integrates bulk RNA-seq and scRNA-seq data by aligning domains via maximum mean discrepancy (MMD) to predict single-cell drug responses. Following this idea, SCAD [23] employs adversarial domain adaptation to predict single-cell drug sensitivity. However, they can only extract domain-invariant features through unsupervised domain adaptation, but failed in determining the decision boundary of sensitivity and resistance in the target domain. Besides, Beyondcell [24] calculates an enrichment score to determine the therapeutic differences among cell populations and generate a prioritized sensitivity-based ranking for drug selection. scRank [25] employs a target-perturbed gene regulatory network to rank drug-responsive cell populations via *in silico* drug perturbations using untreated single-cell transcriptomic data. Nevertheless, they either depend on human-designed scores to rank drugs or prior knowledge to identify cell populations of interest.

In recent years, semi-supervised domain adaptation methods have achieved better performance by leveraging a small number of labeled samples in target domain [26– 30]. Inspired by semi-supervised domain adaptation [31] and adversarial training [32], we introduce SSDA4Drug to transfer drug response from the source domain (bulk RNA-seq) to the target domain (scRNA-seq). By alternately maximizing the conditional entropy of unlabeled target samples with respect to the classifier and minimizing it with respect to the feature encoder, SSDA4Drug successfully reduces the distribution gap between source and target domains, and learns the discriminative features specific to the classification task, which significantly improves the performance in target domain. Remarkably, we found that with only a few labeled target-domain samples, SSDA4Drug effectively overcomes the limitations of unsupervised domain adaptation methods and greatly enhances drug response predictive performance at single-cell level. Our extensive experiments across multiple independent scRNA-seq datasets confirm that SSDA4Drug achieves superior performance compared to five existing methods. Moreover, SSDA4Drug can not only effectively identify the temporally dynamic changes in single-cell drug sensitivity during tumor cell exposure to drugs, but also accurately predict the developmental trend in drug sensitivity along cancer cell evolutionary trajectories. Finally, by exploring the SHapley Additive exPlanations (SHAP) values and integrated gradients of each input gene, we successfully uncover several genes relevant to drug responses.

## Results

### Semi-supervised few-shot transfer learning

SSDA4Drug is a deep learning framework that leverages semi-supervised domain adaptation techniques to translate drug response knowledge from source domain to the target domain (Fig. 1a; Supplementary Fig.7). In this study, we designate bulk RNA-seq data from cell lines as the source domain and scRNA-seq data as the target domain. Within the source domain, each cell line has been assigned a response label for specific drug, whereas only a limited number of cells in the target domain carry such labels. Our model takes inspiration from adversarial perturbations [32] and minimax entropy techniques [31].

**Fig. 1:**
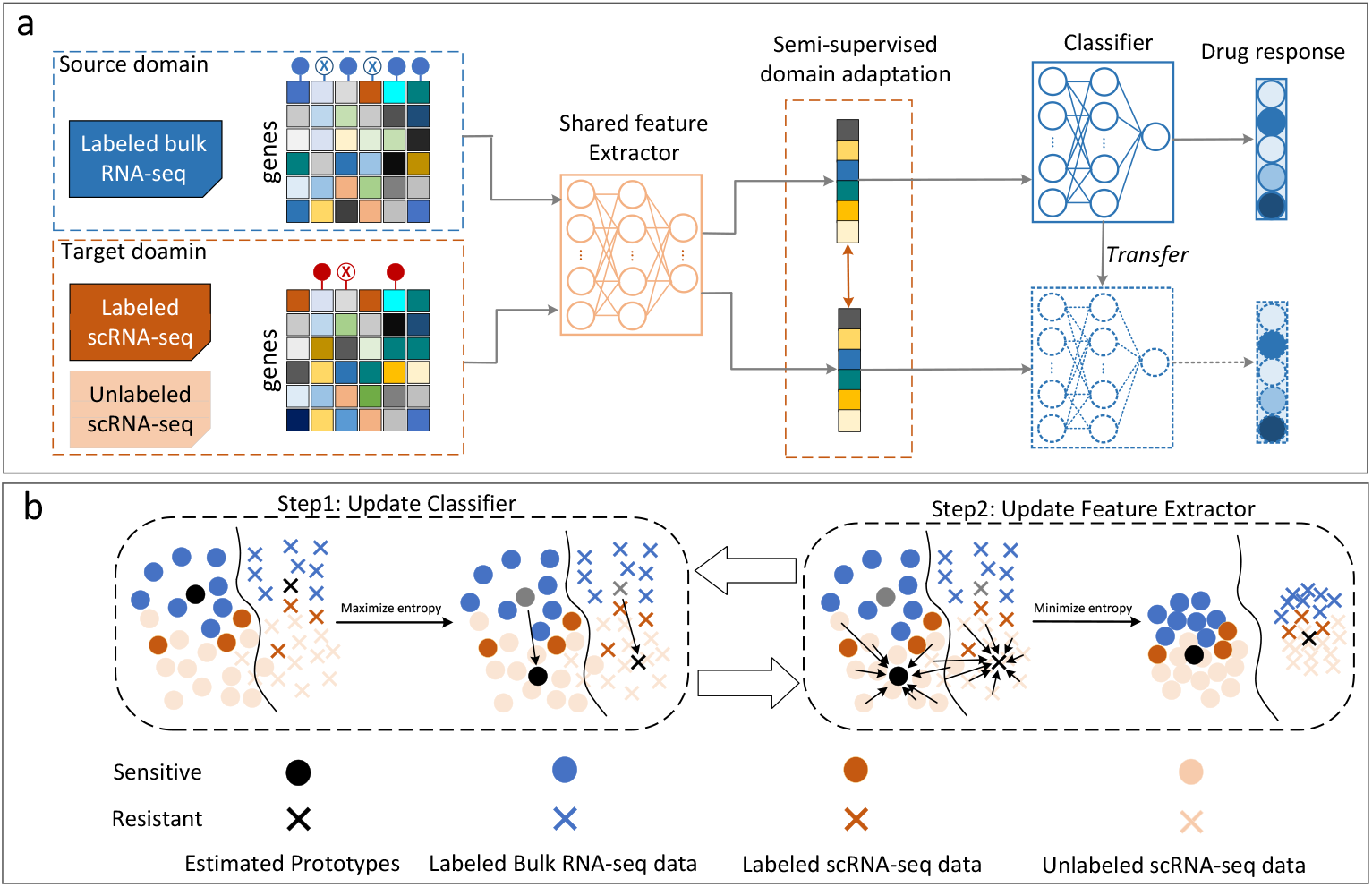
Overview of model architecture and illustration of semi-supervised domain adaptation. (a) Extraction of discriminative and domain-invariant features via semisupervised domain adaptation to predict drug response for target-domain samples. (b) Illustrative diagram of semi-supervised domain adaptation based on minimax entropy. The feature extractor and classifier are trained through adversarial learning to fully exploit unlabeled target samples, thereby deriving better features for classification tasks.

We utilize a shared denoising autoencoder to extract the low-dimensional representations for both source and target domains. A fully-connected network functions as a classifier to determine drug response labels. To effectively exploit the unlabeled target samples for extracting discriminative features, the class prototypes (representatives of each class) are trained to gravitate towards target samples by maximizing the conditional entropy of unlabeled target examples with respect to the classifier, while the feature extractor is updated to minimize the entropy, thereby better clustering them around their respective prototypes (Fig. 1b). By optimizing the minimax loss on the conditional entropy, coupled with adversarial perturbations to bolster the local smoothness, SSDA4Drug not only reduces the distribution gap between domains but also learns discriminative features specific to the classification task.

### SSDA4Drug achieves high performance in predicting single-cell drug responses

We first evaluate the predictive power of our method on single-cell drug responses. The datasets used for performance evaluation included ten single-cell drug response datasets from five distinct cell lines treated with nine chemotherapy and targeted drugs (Table 1). It is important to highlight that these datasets correspond exclusively to the target-domain samples, whereas the source-domain data is derived from the GDSC project, comprising the bulk RNA-seq and drug sensitivity data. Since differentially expressed genes between pre-treatment and post-treatment reflect the substantial effect of drug on cellular context, we selected a set of highly variable genes shared between the source and target domains as model input. Specifically, we identified the highly variable genes using the function of highly variable genes in Scanpy [33], and then selected genes involved in protein–protein interactions from a prior knowledge graph that has compiled the most informative genes relevant to drug sensitivity [34]. Meanwhile, to balance the proportion of sensitive and resistant cell lines in the source domain, we leveraged a weighted-sampling algorithm [35] to generate samples in the minority class, by increasing the likelihood of minority categories being sampled assigning higher weights to minority categories and lower weights to majority categories.

**Table 1:** Details of the single-cell drug response datasets for performance evaluation [23].

| Drug | Dataset | Cell line | #res. | #sen. | #gene | Cancer type |
| --- | --- | --- | --- | --- | --- | --- |
| Etoposide | GSE149215 | PC9 | 764 | 629 | 2668 | Lung cancer |
| PLX4720 | GSE108383 | 451Lu | 81 | 74 | 2856 | Melanoma |
| PLX4720 | GSE108383 | A375 | 46 | 63 | 2873 | Melanoma |
| Vorinostat | CCLE | JUH006 | 33 | 33 | 2933 | Oral squamous cell carcinomas |
| Gefitinib | CCLE | JUH006 | 33 | 33 | 2933 | Oral squamous cell carcinomas |
| AR-42 | CCLE | JUH006 | 33 | 33 | 2933 | Oral squamous cell carcinomas |
| NVP-TAE684 | CCLE | SCC47 | 60 | 60 | 2876 | Oral squamous cell carcinomas |
| Afatinib | CCLE | SCC47 | 60 | 60 | 2876 | Oral squamous cell carcinomas |
| Sorafenib | CCLE | SCC47 | 60 | 60 | 2876 | Oral squamous cell carcinomas |
| Cetuximab | CCLE | SCC47 | 60 | 60 | 2876 | Oral squamous cell carcinomas |

For each dataset listed in Table 1, we built a model and evaluated its performance separately. It is worth noting that only 3 cells from each category were randomly selected as labeled target-domain samples, while the remaining cells served as unlabeled samples for semi-supervised training and performance evaluation. To avoid deviation induced by randomness, we repeated the process 50 times and calculated the mean AUC and mean AUPR as performance metrics. To benchmark our method, we compared our method with four other methods. First, we established a baseline model that is trained on the GDSC bulk RNA-seq data and then directly used to predict drug responses of single cells. Moreover, we conducted performance comparison with three current state-of-the-art methods, including scDEAL [22], CODE-AE [9] and SCAD [23]. We evaluated two variants of scDEAL, one with pretraining and one without. The experimental results indicate that our method consistently outperforms other approaches in nearly all ten drug–cell type combinations (Fig. 2a). Except for Etoposide, our method achieved AUC values exceeding 0.8 across all other drug–cell type combinations. Although CODE-AE and SCAD match our method’s performance for certain drugs, they exhibit lower stability overall and perform considerably worse on others. Meanwhile, scDeal with pretraining displays marked instability, with outcomes heavily influenced by random data initialization.

**Fig. 2:**
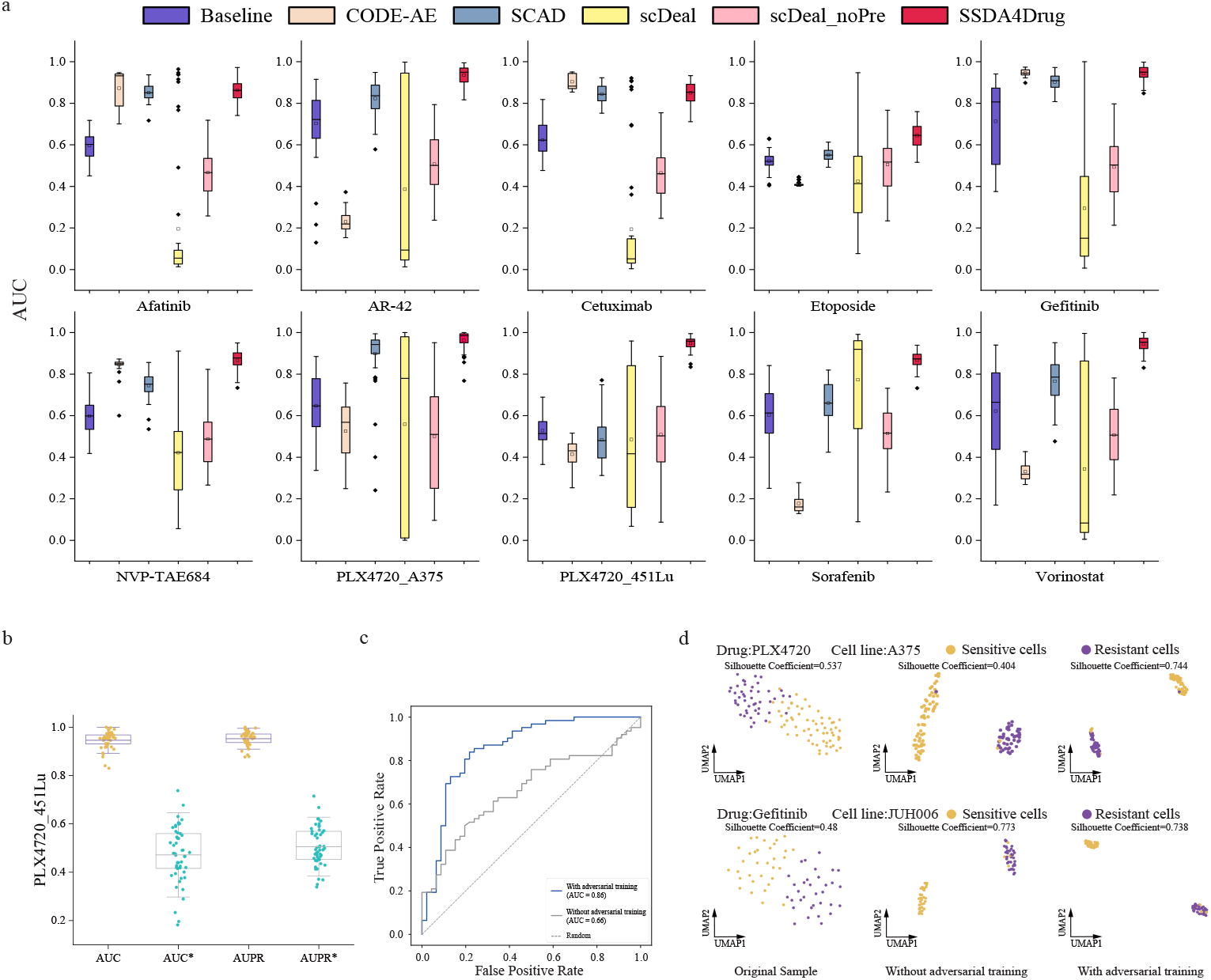
Evaluation of SSDA4Drug performance in predicting single-cell drug responses. (a) Performance comparison of our method with five other existing methods across ten single-cell drug response datasets (b) Boxplots of the AUC and AUPR values of SSDA4Drug trained without any melanoma cell lines in source domain for predicting 451Lu cell response to PLX4720 treatment. (c) ROC curves and AUC values of SSDA4Drug trained without any melanoma cell lines in source domain for predicting A375 cell response to PLX4720 treatment. (d) UMAP plots of the primary expression profiles and embeddings obtained by SSDA4Drug with and without adversarial training for the A375 and JUH006 cells treated by PLX4720 and Gefitinib, respectively.

To assess our model’s generalizability, we evaluated its performance across two different melanoma cell lines (451Lu and A375) treated with the same drug (PLX4720). In particular, we excluded all melanoma cell lines in the source domain from GDSC bulk RNA-seq dataset. For the target domain, we randomly selected only 3 PLX4720-treated 451Lu cells as labeled samples to train the model, which was then used to predict the drug response of the remaining 451Lu cells. Our model achieved an AUC and AUPR of approximately 0.95 for 451Lu single-cell responses prediction (Fig. 2b). Notably, adversarial training significantly improved the performance. By excluding all cell lines of a specific cancer type from the source domain, this experiment poses a stringent challenge for generalizability assessment. The robust performance of our model highlights its ability to learn drug-response features from cell lines with similar expression profiles, thereby enhancing its generalizability. Furthermore, we conducted a cross-cell-type prediction task to evaluate the model’s capabilities by applying the trained model to predict the response of A375 cells to PLX4720. This task provided an even more rigorous test of the model’s generalization capability. With adversarial training, our model achieved a substantially high performance (AUC=0.86) (Fig. 2c). These experiments confirm the strong generalizability of our method, demonstrating its effectiveness in predicting single-cell drug responses even when trained on data from distinct cancer types.

To validate the beneficial effects of adversarial training on model robustness, we applied Uniform Manifold Approximation and Projection (UMAP) dimensionality reduction [36] to visualize the latent representations of A375 and JUH006 cells treated with PLX4720 and Gefitinib, respectively. The UMAP plots showed that adversarial training led to tighter clusters of intra-class samples from the target domain in the latent space, while inter-class samples were more widely separated (Fig. 2d and Supplementary Fig.1). Also, it can be found that the silhouette coefficients for clusters derived from primary expression profiles are relatively low. In contrast, when using the features extracted by our model, the silhouette coefficients markedly increase, with most values exceeding 0.5. This observation indicated that drug-resistant and sensitive cells are well separated in the latent space, thereby leading to high predictive performance.

### Few-shot in target domain brings great performance enhancement

For systematic evaluation of few-shot impact on performance, we gradually increased the number of labeled target samples included in the training set. Using the datasets in Table 1, we employed distinct increments in labeled target samples, due to differing numbers of single cells across various drugs. For Gefitinib and Sorafenib, we progressively increased the labeled cells from 1 to 4. For Etoposide and PLX4720-451Lu, we used 1, 2, 5, and 10 labeled cells for model training, respectively. In each few-shot task, the designated number of labeled target samples was randomly selected for the training set, with the remainder used to assess model performance. To mitigate randomness-induced deviations, we repeated each few-shot task 50 times.

The results revealed a significant enhancement in performance for all four drugs as the number of labeled target samples increased (Fig. 3 and Supplementary Fig.2). Specifically, for Gefitinib and Sorafenib, the mean AUC values started at 0.912 and 0.782 for 1-shot scenario and increased by 4.6% and 11.2% at 4-shot scenario, respectively (Fig. 3a). In the case of Etoposide and PLX4720, their performance at the

**Fig. 3:**
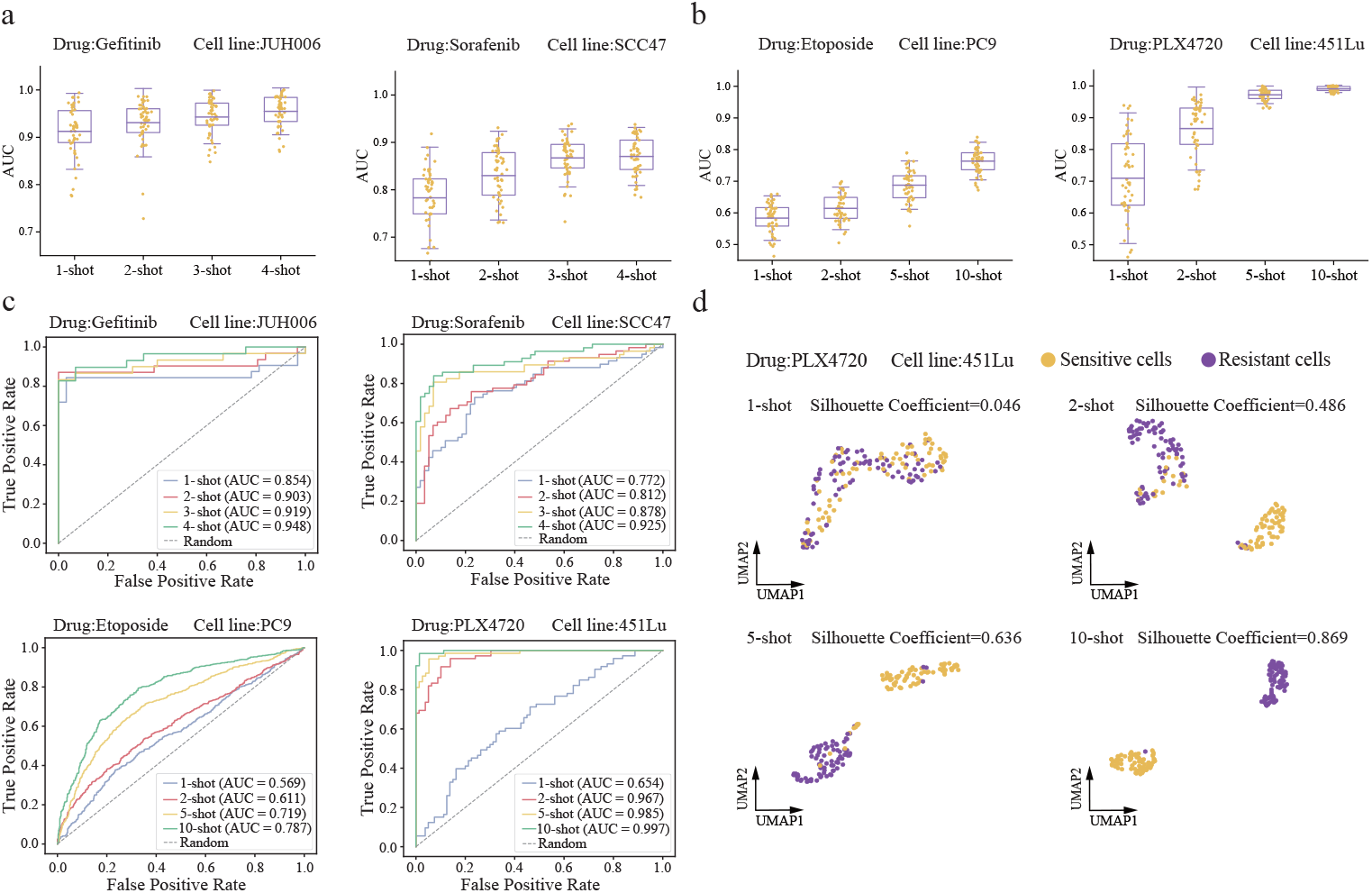
Performance enhancement of SSDA4Drug brought by increasing number of shots in target domain. (a) Boxplot of AUC values on Gefitinib-JUH006 and Sorafenib-SCC47 scRNA-seq datasets by SSDA4Drug trained using 1, 2, 3, and 4 labeled targetdomain shots. (b) Boxplots of AUC values on Etoposide-PC9 and PLX4720-451Lu scRNA-seq datasets, obtained by SSDA4Drug trained using 1, 2, 5, and 10 labeled target-domain shots. (c) ROC curves and AUC values on four different scRNA-seq datasets achieved by SSDA4Drug trained with varying number of target-domain shots, respectively. (d) UMAP plots of the latent representations for PLX4720-451Lu scRNA-seq data achieved by SSDA4Drug trained with different number of target-domain shots.

10-shot scenario gained a substantial boost of 31% and 39% in mean AUC values compared to the 1-shot scenario, respectively (Fig. 3b, Supplementary Data 2). A close-up view of the ROC curves and AUC values revealed the significant performance improvement brought about by the increase in shots across the four drug-induced single-cell response datasets (Fig. 3c). Notably, the variance in AUC for PLX4720-451Lu dataset decreased greatly with the increasing number of shots, indicating improved model reliability.

To visualize these effects, we utilized UMAP plots to depict the embeddings of target samples from the PLX4720-451Lu dataset, which were derived from models trained with varying shot numbers (Fig. 3d). Evidently, as the number of shots increased, the intra-class single-cell embeddings clustered closer together, while the inter-class samples moved further apart. These findings not only confirmed the benefits of few-shot learning in predicting single-cell drug responses, but also demonstrated that our model’s discriminatory power and reliability boosted with the increasing number of few-shot samples.

### SSDA4Drug captures dynamic state transition toward drug resistance

Previous studies have shown that cancer cells often evolve rapidly in response to therapeutic pressure, and eventually develop resistance to specific drugs [37]. To further validate the performance of SSDA4Drug, we assessed its ability to predict temporally dynamic changes of drug response for individual cells during persistent chemotherapy drug exposure. For this purpose, we used the dataset published by Ben-David et al. [38], which encompassed 7,440 single-cell clones derived from MCF7 cell line. These cells were exposed to 500nM Bortezomib, a 26S proteasome inhibitor, for 2 days. After filtering out low-quality data (for detail see Method), 3,740 cells were retained for subsequent analysis. They underwent single-cell sequencing at various time points: before treatment (t0, *n*=160), 12 hours after drug exposure (t12, *n*=994), 48 hours after exposure (t48, *n*=1623), and 96 hours after drug washout and recovery (t96, *n*=963) (Fig. 4a). The UMAP plots revealed distinct clustering of the transcriptional profiles of these cells into four groups, implying rapid transcriptional state transitions induced by drug exposure.

**Fig. 4:**
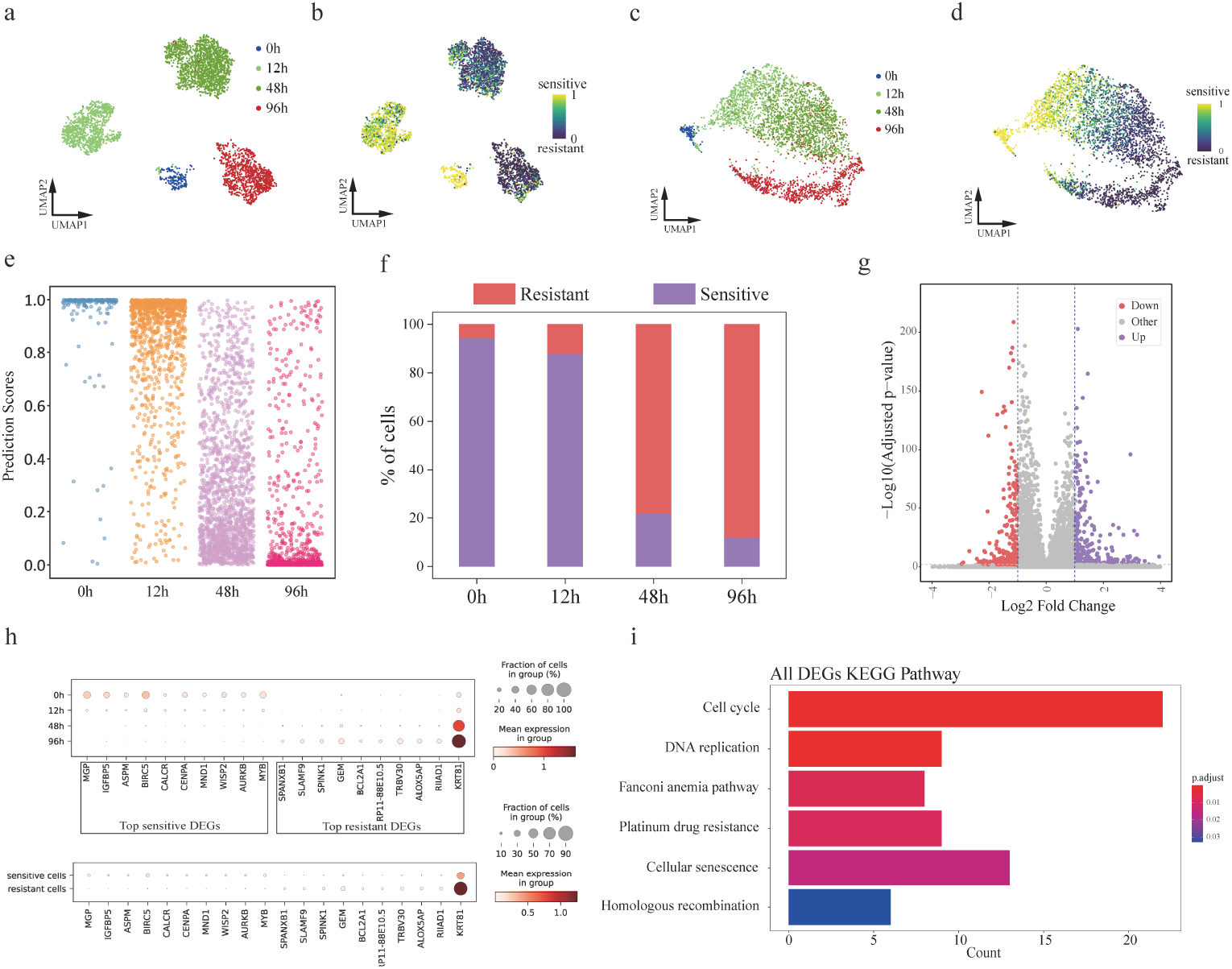
SSDA4Drug accurately predict the temporal dynamics of 3,740 MCF7 cells in response to Bortezomib at four time points. (a-b) UMAP plots of the scRNA-seq data all cells exposed to Bortezomib colored by time points and predicted sensitivity scores, respectively. (c-d) UMAP plots of the latent representations obtained by SSDA4Drug for all cells colored by time points and SSDA4Drug predicted scores, respectively. (e) Scatter plots of SSDA4Drug’s predicted scores for all cells at four time point. (f) Percentage of sensitive and resistant cells predicted by SSDA4Drug at each time point. (g) Volcano plot for differential expression analysis between sensitive and resistant cell groups predicted by SSDA4Drug. (h) Upper and lower plots show the cell fraction and normalized expression levels of top 10 differentially expressed genes across four time points and predicted cell groups, respectively. (i) KEGG enrichment analysis for the differentially expressed genes between sensitive and resistant cells predicted by SSDA4Drug.

We tested whether SSDA4Drug could accurately predict the temporal variations of MCF7 cells in response to Bortezomib based on the scRNA-seq data. To this end, we assigned 20% of the cells at t0 (*n*=32) as sensitive and 20% of the cells at t96 (*n*=193) as resistant. Then, we randomly selected only 3 cells from each category as labeled target samples, with the remaining serving as unlabeled target samples for model training. All cells were used to verify SSDA4Drug’s ability to capture the temporal changes in drug efficacy at different time points. We analyzed the predicted scores of individual cells, and revealed that almost all MCF7 cells were sensitive to Bortezomib before treatment. However, as drug exposure duration increased, the number of sensitive cells gradually decreased while the number of resistant cells increased. At 96 hours post-treatment, the cell population contained the highest proportion of resistant cells (Fig. 4b,e). The UMAP plots showed that the learned embeddings of the cells diverged gradually from sensitivity to resistance induced by drug treatments (Fig. 4c,d). Our results were highly consistent with the findings of a previous study [39].

We were intrigued by the molecular mechanism that underlies the transition of cellular state from sensitive to resistant. For this purpose, we categorized all cells as either resistant or sensitive based on the predicted scores using a threshold of 0.5. This categorization revealed a gradual increase in the proportion of resistant cells over time (Fig. 4f). Subsequently, we conducted differential expression analysis (logFC *>*1 and p-adj *<*0.05) between the two groups of cells with assigned response labels, and identified 385 drug-sensitive genes and 255 drug-resistant genes (Fig. 4g). Among them, we selected the top 10 overexpressed genes from both sensitive and resistant DEGs, and visualized their respective cell proportions and gene expression levels at each time point (Fig. 4h). Two distinct gene expression patterns emerged when comparing t0 and t12 to t48 and t96. Among the top 10 drug-resistant DEGs, a few genes have been reported to be associated with drug resistance(Supplementary Table 2). For instance, BCL2A1 has been shown that its high expression in tumor samples correlates with in vivo chemoresistance [40]. SPINK1 overexpression correlates with Hepatocellular Carcinoma treatment Resistance [41]. Finally, we performed KEGG functional enrichment analysis on the differentially expressed genes and found that the overexpressed genes were enriched in cell proliferation-related signaling pathways, such as cell cycle and DNA replication (Fig. 4i). These experimental results demonstrated the capacity of our method to predict the dynamics of cellular responses to drug treatment across multiple time points.

### SSDA4Drug identifies reversible drug response states of lung cancer cells

To further evaluate whether SSDA4Drug can identify reversible drug response states of cancer cells, we applied it to the dataset published by Aissa et al. [42]. This dataset comprised the Drop-seq data from lung cancer PC9 cells cultured in vitro, including untreated cells, Erlotinib-treated cells, and cells underwent a drug holiday (Fig. 5a). After filtering out low-quality data, 3,053 cells were retained for analysis. Specifically, these cells covered various time points during Erlotinib treatment: before treatment (D0, *n*=765), after 1 day (D1, *n*=236), 2 days (D2, *n*=148), 4 days (D4, *n*=100), 9 days (D9, *n*=228), and 11 days (D11, *n*=143) of drug exposure, as well as the cells followed by re-treatment for 2 days after a 6-day drug holiday (D19ERL, *n*=726) and cells followed an 8-day drug holiday (D19DMSO, *n*=707). The number of living cells decreased from D1 to D4 during drug exposure and then increased thereafter, indicating a transition from Erlotinib sensitivity to resistance in PC9 cells (Supplementary Fig.3). The cell subpopulation that has developed resistance survived and proliferated, thereby leading to growth in cell numbers even during drug exposure. In contrast, the D19DMSO cells regained sensitivity to Erlotinib to a large extent, implying that drug holiday successfully reversed acquired drug resistance [42]. In fact, previous studies have suggested the possibility of temporary reversal of drug resistance, exemplified by the “drug holiday” phenomenon observed during clinical therapy by EGFR inhibitors [43].

**Fig. 5:**
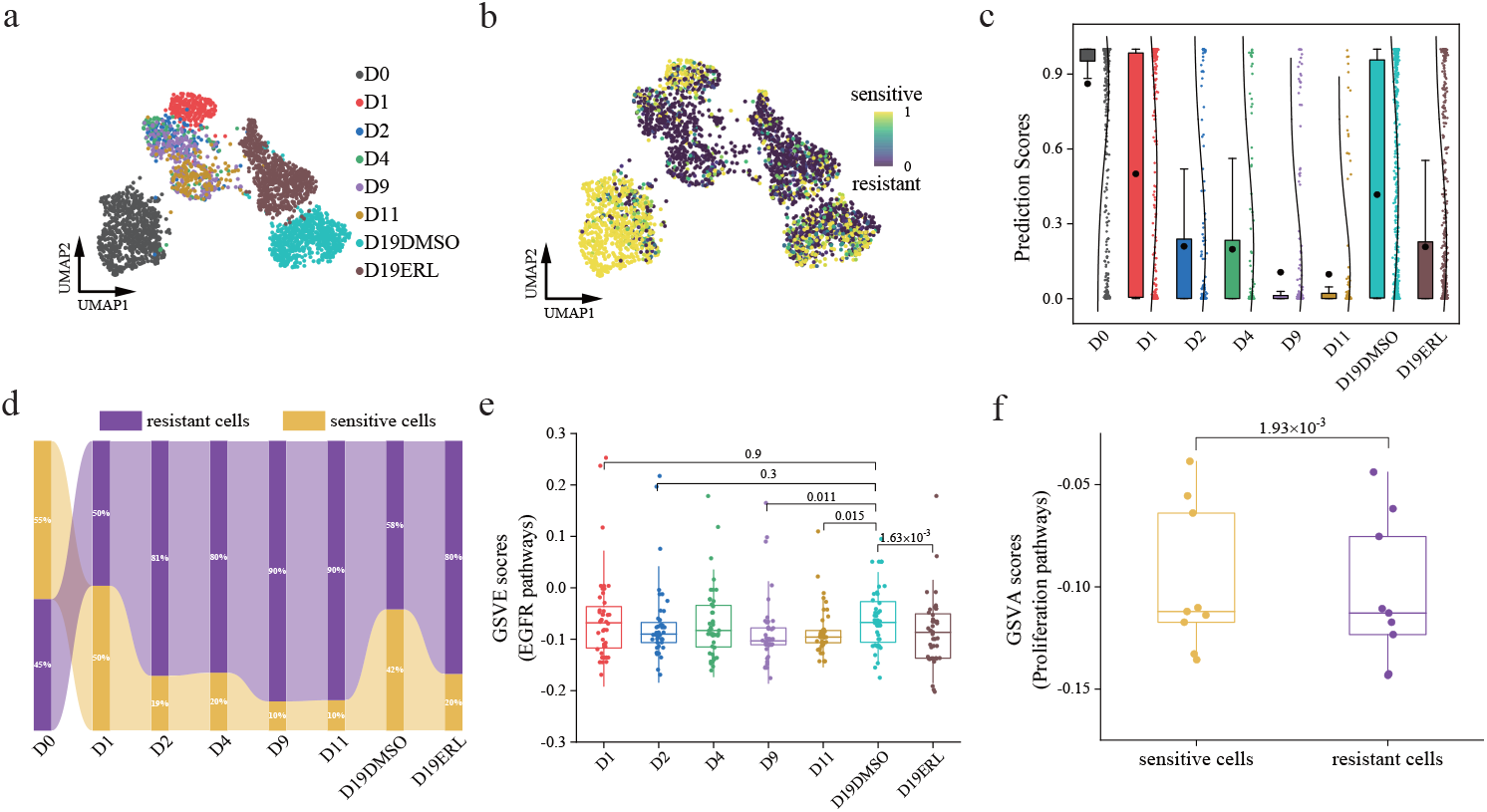
SSDA4Drug successfully identified reversible drug response states of PC9 cells subjected to Erlotinib treatment. (a-b) UMAP plot of scRNA-seq data of PC9 cells exposed to Erlotinib, colored by eight time points and predicted sensitivity scores, respectively. (c) Boxplots of SSDA4Drug’s predicted scores for PC9 cells at eight time points. (d) Area plot of the percentage of sensitive and resistant cells predicted by SSDA4Drug at each time point. (e) Boxplots of GSVA scores regarding the activity levels of EGFR-related pathways. P-values were obtained from the paired-sample t-test. (f) Boxplot of GSVA scores regarding the activity levels of proliferation-related pathways in the sensitive and resistant cell groups. P-values were obtained from the paired-sample t-test.

Our objective was to evaluate SSDA4Drug’s predictive ability regarding reversible drug response states and explore the underlying biological mechanisms. We labeled the cells before treatment (D0) as sensitive and the cells received 9 days (D9) and 11 days (D11) treatments as resistant. These cells were used as labeled target-domain samples to train the model, which was then used to predict the drug response states of the cells at other time points. Consistent with the actual observations, the numbers of sensitive cells predicted by SSDA4Drug declined from D1 to D11 during drug exposure (Fig. 5b,c), while the number of cells predicted to be resistant increased. As expected, the predicted proportion of sensitive cells in D19DMSO significantly increased after the drug holiday (Supplementary Fig.4). According to the predicted scores (higher score means higher sensitivity), we categorized individual cells as either sensitive or resistant using a predicted value threshold of 0.5. As a result, the proportion of sensitive cells was found to gradually decrease but increased again after the drug holiday (Fig. 5d). Specifically, D19DMSO cells showed a 22.8% increase in sensitive cells compared to D2 cells, and a 22.3% increase compared to D19REL cells re-treated after the drug holiday. These findings demonstrate a high degree of consistency between our predictions and the actual situation.

Considering that Erlotinib is an inhibitor of EGFR pathway, we employed the GSVA tool [44] to assess the activity levels of EGFR-related signaling pathways (Supplementary Data 5) in each cell. The mean GSVA scores across all cells at various time points served as indicators of pathway activity levels. The results revealed a correlation between the GSVA scores for EGFR pathway and the proportion of drug-sensitive cells, elucidating the dynamic shifts in cellular sensitivity to the drug treatment (Fig. 5c,e).

Additionally, drug-sensitive cells exhibited higher GSVA scores in cell proliferation-related signaling pathways compared to drug-resistant cells (Fig. 5f). The findings were consistent with previous studies indicating that actively proliferating cells exhibit high sensitivity to specific anticancer drugs [8]. These collective insights demonstrated that our method actually captures pharmacogenomic features related to drug response from expression profiles, rather than just memorizing the samples, thereby achieving strong generalizability.

### SSDA4Drug predicts development of drug resistance along with evolutionary trajectory

We seek to explore the evolutionary trajectory of cancer cells and its intricate linkage to the development of drug resistance. When subjected to the selective pressure of drug therapy, cancer cells undergo genetic alterations and phenotypic shifts, which enable them to withstand the lethal effects induced by therapeutic agents. This adaptation poses significant challenges to the treatment efficacy. Therefore, it is important to identify the pivot point where cancer cells transition from being drug-sensitive to drug-resistant during their evolutionary journey. This motivation led us to explore the correlation between the drug response states predicted by SSDA4Drug and the evolutionary trajectory of tumor cells.

To this end, we conducted analysis on the GEO dataset (accession number GSE117872) [45]. This dataset comprises the scRNA-seq data of 1,302 cells collected from three patients with oral squamous cell carcinomas (OSCC). Each cell in the dataset has been labeled as either sensitive or resistant to Cisplatin. We randomly selected 50 cells as labeled target samples to train the model, and subsequently assessed its performance on the remaining cells. The results showed that 88% of the cells were correctly predicted, with an AUC score of 0.927 and an AUPR score of 0.978. Among them, 98% of the predicted resistant cells and 84.6% of the predicted sensitive cells matched their ground-truth labels (Fig. 6a). Next, we performed pseudotime analysis on the scRNA-seq data using Monocle3 [46]. The result reflects the inferred evolutionary trajectory based on the transcriptional profiles of these cancer cells (Fig. 6b). By depicting the actual drug response state of the cells along the trajectory, we could clearly observe the progression of cancer cells from the sensitive state to resistant state (Fig. 6c). As expected, our comparative analysis between the predicted drug resistance progression and the ground-truth data confirmed that the development of drug resistance is tightly coupled with the evolutionary trajectory of cancer cells (Fig. 6c,d). These findings suggested that our method successfully identifies the transcriptional patterns of cancer cells shifting from sensitivity to resistance along the evolutionary trajectory.

**Fig. 6:**
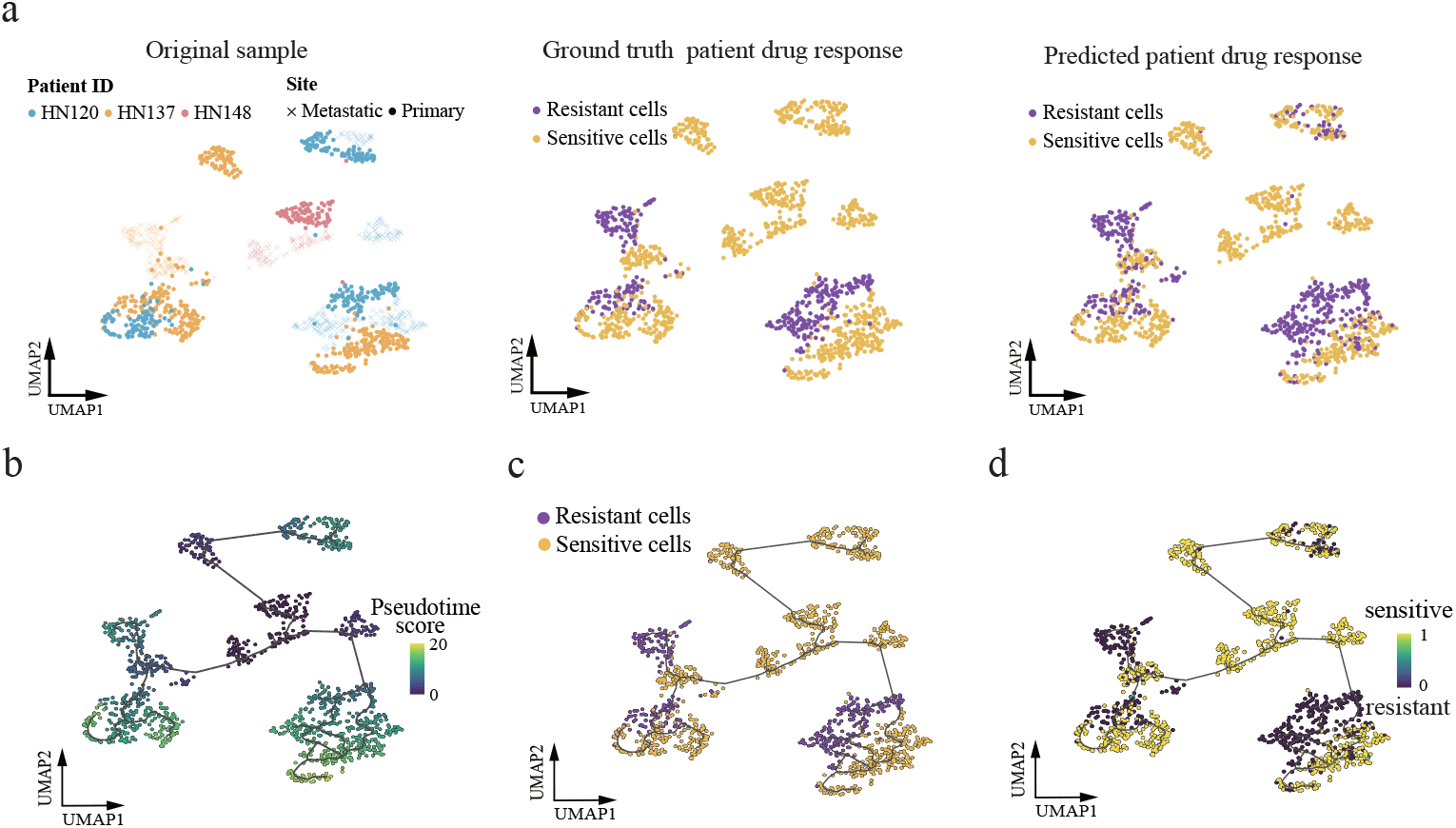
SSDA4Drug predicted drug resistance progression highly coupled with the evolutionary trajectory of cancer cells when subjected to the selective pressure of Cisplatin treatment. (a) UMAP plots of the scRNA-seq data of OSCC cells colored by sample sources, ground-truth drug response labels, binarized drug response labels based on predicted scores, respectively. (b-d) Pseudotime trajectory of the OSCC cells in which the cell was colored by pseudotime scores, ground-truth drug response labels, and predicted drug response probability, respectively.

### SSDA4Drug uncovers key genes relevant to drug response

We were interested in exploring the ability of SSDA4Drug to uncover critical genes determining the single-cell drug response. We applied it to another single-cell drug response dataset (GEO accession number GSE140440) [47]. This dataset comprises the scRNA-seq profiles of 324 cells derived from two prostate cancer cell lines, PC3 and DU145. Each cell has been assigned a label either sensitive or resistant to Docetaxel (Fig. 7a). We randomly selected three cells as labeled target-domain samples to train the model. For performance evaluation on this dataset, we found that 96% of the cells were correctly predicted for their sensitivity or resistance to Docetaxel, with an AUC value of 0.97. Based on the predicted scores, we classified each cell as either resistant or sensitive using the threshold of 0.5, and found that the predicted drug responses aligned almost perfectly with the ground-truth drug responses.

**Fig. 7:**
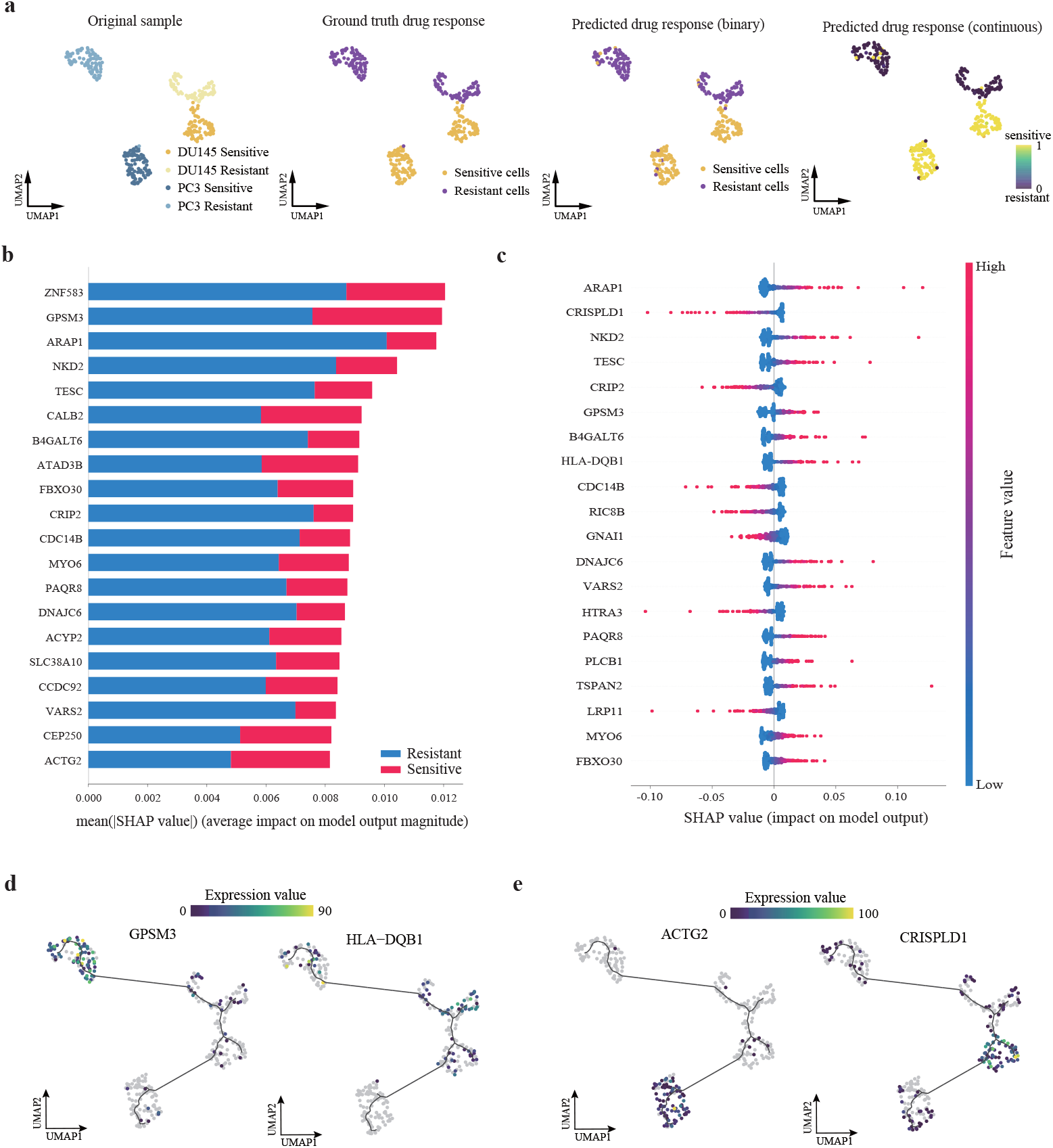
Discovery of key genes involved in the drug response of PCa cells to Docetaxel treatment. (a) From left to right: UMAPS plots of cells colored by tissue sources, ground-truth labels, binarized drug response labels and predicted scores, respectively.(b) Mean absolute SHAP values of top 20 genes contributing significantly to the drug response state of PCa cells. (c) SHAP beeswarm plot of top 20 genes contributing to drug resistance in PCa cells. (d) Pseudotime trajectory plots for two representative drug-resistant genes, GPSM3 and HLA-DQB1, colored by their respective gene expression levels. (e) Pseudotime trajectory plots of two representative drug-sensitive genes, ACTG2 and CRISPLD1, colored by their respective gene expression levels.

Following proof of performance, we employed the SHAP values [48] to uncover the key genes involved in drug response. Therefore, we calculated the mean absolute SHAP values for each gene in sensitive and resistant cell groups, representing their average impact on the drug response predictions (Fig. 7b). To further illustrate the contributions of the genes to resistance and sensitivity, we generated beeswarm plots for the resistant cell group (Fig. 7c) and the sensitive cell group (Supplementary Fig.6). By examining two representative Docetaxel-resistant genes, GPSM3 and HLA-DQB1 (Fig. 7d), we observed their increased expression levels closely coupled with the transition of cells towards drug-resistant phenotype (Fig. 7a,d). Conversely, two representative Docetaxel-sensitive genes, ACTG2 and CRISPLD1, demonstrated an opposite expression pattern (Fig. 7e). Previous studies have reported GPSM3 as a potential immunotherapy target in low-grade Gliomas [49], while ACTG2 has been shown to control the growth of small intestinal neuroendocrine tumor (SI-NET) cells [50].

In addition, we leveraged the integrated gradients (IG) [51] to uncover key genes. To this end, we computed the integrated gradient score for each gene in individual cells, and then ran differential expression analysis (sensitive vs resistant) based on their IG values (Supplementary Data 4). We visualized the top 10 genes with the highest IG values in both resistant and sensitive cells (Supplementary Fig.5), from which we found distinct patterns between the two cell populations. These findings demonstrated our model’s interpretability for identifying the genes that contributed most significantly to drug responses.

## Discussion

Tumor heterogeneity often leads to significant differences in the response to the same drug treatment among different cancer cells. The advancement of single-cell sequencing technology has opened up new avenues to identify pre-existing and acquired drug-resistant tumor cell subpopulations. However, the currently available single-cell drug response data covers only a few cancer types and drugs, due to cost and technical limitations. Therefore, the computational methods with strong predictive power are a practical solution to determine single-cell drug response. However, the limited availability of scRNA-seq data with drug response information makes it challenging to develop a predictive model that relies solely on single-cell data. In this paper, we utilized semi-supervised domain adaptation to transfer the pharmacogenomic knowledge from bulk-level to single-cell level, establishing a more effective method for building a predictive model with superior performance. The experimental results demonstrated that our method possessed sufficient generalizability and notably outperformed previous methods based on unsupervised domain adaptation.

In our evaluation experiments, we observed that predicting drug responses for certain cell types poses significant challenges. Specifically, for PC9 cells treated with Etoposide, the average AUC attained by competing methods consistently falls below 0.6 (Fig. 2a). The UMAP plots of PC9 cells’ primary expression profiles (Supplementary Fig. 1) indicate that sensitive and resistant cells exhibit highly overlapping patterns, making them difficult to distinguish and likely contributing to the suboptimal predictive performance of these methods. Our method incorporates a small number of labeled target-domain samples to guide the encoder toward learning discriminative features of PC9 cells. Additionally, adversarial training further enhanced its ability in feature extraction, as demonstrated by the UMAP visualizations that show a significant separation between sensitive and resistant cells in the latent space (Supplementary Fig. 1). As a result, our method achieved a markedly higher average AUC of 0.71, substantially outperforming all competing methods. Also, in our investigation of the reversible drug response states of PC9 cells exposed to Erlotinib, the wet-lab data revealed that the proportion of resistant cells progressively increases with prolonged drug exposure (Fig. 5a, Supplementary Fig.3). However, close examination of the single-cell expression profiles revealed that, at certain time points, sensitive and resistant cells remain highly intermixed (Fig. 5b). Particularly, some resistant cells revert to sensitive state after a drug holiday. Predicting such dynamic shifts in cellular states is particularly challenging. Our model, trained using only the data of one or two time points, successfully predicted the dynamic drug response trends across multiple other time points (Fig. 5d). In contrast, competing methods have not underwent performance evaluation under similarly stringent conditions yet.

One limitation of our current model lie in its drug-specific design, which restricts its applicability to broader scenarios. To address this, we could incorporate a pre-trained drug molecular encoder into our framework, drawing inspiration from chemCPA [52] and cycleCDR [53]. These models have demonstrated their capability to predict pandrug cellular transcriptional responses upon various drug perturbations. Integration of a drug molecular encoder would enable the extension of our model into a robust pandrug framework. On the other hand, the limited availability of single-cell response data for specific drugs currently constrains our model to a target domain comprising only a single cancer type. Consequently, the present implementation is tailored for single-cell drug response prediction within this narrow context. However, our model architecture is inherently flexible and can be easily adapted to a pan-cancer framework without any structural modifications, provided that the input dataset includes scRNA-seq profiles from multiple cancer types. We believe that the growing availability of single-cell drug response datasets will effectively address this limitation. For instance, the cDrugAtlas [54]and DRMref [55] databases have already curated extensive single-cell response data across multiple cancer types treated with identical drugs.

Finally, this study primarily used the gene expression profiles for feature extraction. Nonetheless, our framework’s flexibility allows for the integration of multi-omics data, including genomic mutations, copy number variations, and epigenetic information. These datasets offer promising opportunities for training a multi-modal model and expanding the scope of our approach.

## Conclusion

In this paper, we proposed a semi-supervised deep transfer learning method for predicting drug responses of individual tumor cells. We have demonstrated that incorporating a few labeled single-cell samples into the domain adaptation not only aligns the distribution of source and target domains but also learns a better classification decision boundary. Our extensive experiments have shown that our model outperformed current state-of-the-art methods [23], but also captures the dynamic changes in drug sensitivity of tumor cell populations during drug exposure, and reversals of response states induced by drug holidays. Moreover, we explored model interpretability to uncover key genes that play an important role in cellular responses to drug treatments.

## Materials and Methods

### Data source and preprocessing

#### Bulk drug sensitivity

The bulk drug sensitivity data were retrieved from the Genomics of Drug Sensitivity in Cancer (GDSC) database [3, 4], which provided drug sensitivities of 1074 cancer cell lines upon 226 drugs. Drug sensitivity was evaluated using half maximal inhibitory concentration (IC50) and area under the dose-response curve (AUC) measurements. We assigned drug response label to each cell line, by using the method similar to CODE-AE [9]. For a drug of interest, we ranked the IC50 values of all cell lines tested against this drug, and then categorized them as sensitive or resistant. The binarization threshold was set as the average IC50 value across all cell lines tested against the drug. The sensitive cell lines were labeled as 1, while the resistant ones were labeled as 0. Together with the drug sensitivity data, we retrieved the matched bulk RNA-seq data (RMA-normalized basal expression profiles) of the cell lines from GDSC.

#### Single-cell drug response

For target-domain data, we collected the scRNA-seq data where cells were derived from different tissue sources, including cell lines, patient-derived xenografts (PDX) models, and patient tumors. The scRNA-seq data and single-cell drug response labels were obtained from the National Center for Biotechnology Information’s (NCBI) Gene Expression Omnibus (GEO). Quality control and preprocessing of the scRNA-seq data were performed using the Python package SCANPY [33]. Specifically, to eliminate lowly expressed genes that may result from noise or background signals, we filtered out cells with fewer than 200 detected genes and genes detected in fewer than 3 cells. To identify and exclude cells that may be contaminated or in an unhealthy state, we also filtered out cell samples with a mitochondrial gene expression ratio exceeding 5%.

#### Overview of SSDA4Drug model

Given the abundant drug response labels in the source domain (cell lines) and the limited number of labels in the target domain (single cells), we proposed the SSDA4Drug model for single-cell drug response prediction. Inspired by minimax entropy [31], our model integrates the semi-supervised domain adaptation and adversarial training techniques, aiming to learn robust decision boundaries for classification tasks and effectively align domain distributions simultaneously (Fig. 1a).

Our model consists of two components: an autoencoder and a classifier. We define the feature extractor ℱ as the encoder plus the network layers except for the last linear layer of the classifier, while the last linear layer of the classifier as 𝒞 (Supplementary Fig.7). We train ℱ and 𝒞 using labeled source-domain samples and a small number of labeled target-domain samples, and evaluate the prototypes of the sensitive and resistant cells. Since the classifier is primarily trained on labeled samples, the estimated prototypes may be biased towards the source samples due to scarcity of labeled target samples. Let the conditional entropy represent the similarity between the estimated prototypes and target features, we firstly train 𝒞 to maximize the entropy between the prototypes and the unlabeled target features. This would pull the prototypes toward the center of the unlabeled target features. Next, we ℱ train to minimize the entropy so that the unlabeled target features are clustered around the prototypes. To summarize, the minimax entropy process is actually the adversarial learning between ℱ and 𝒞. Repeating such entropy maximization and entropy minimization process yields discriminative features. Meanwhile, we utilize adversarial perturbations to enhance the local smoothness of the model. Adversarial samples regarding data points near the decision boundary could clarify the decision boundaries between drug-sensitive and drug-resistant cells.

### Denoising autoencoder for feature extraction

We utilize a common denoising autoencoder to extract a low-dimensional representation for both the source and target domains, taking as input the expression profiles of labeled and unlabeled samples. Let 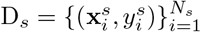 represent *N*_*s*_ labeled source samples, where **x**_*i*_ represents the expression profile and *y*_*i*_ = {1, 0} represents the corresponding label (sensitive or resistant) of *i*-th example upon specific drug treatment.

Similarly, 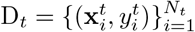 represents *N*_*t*_ labeled target samples, and 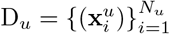 represents *N*_*u*_ unlabeled target samples (*N*_*t*_ ≪ *N*_*u*_).

To learn robust features, we introduce random noise generated from a binomial distribution to each gene expression value, so that we get the noisy expression profile **x**′. The encoder maps the noisy gene expression profiles to a low-dimensional latent representation *z*, while the decoder strives to reconstruct the original noise-free data from this latent representation. To minimize the reconstruction error, we employ the mean squared error (MSE) loss function as below

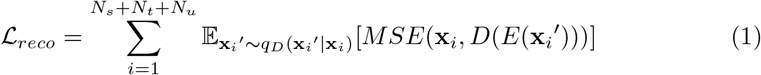

where *q*_*D*_(**x**_*i*_′|**x**_*i*_) represents the probability distribution from which the noise drawn.

### Drug response predictor

Suppose that the final linear layer of the classifier, i.e. 𝒞 consists of weight vectors W_*C*_ *∈* R^*d×c*^, which maps the features ℱ (**x**) extracted by previous layers to binary class probabilities. Since previous studies have demonstrated that normalization can enhance model stability and performance, we normalize the features and feed them into the last linear layer 𝒞, followed by a softmax layer with a temperature parameter *T*. As a result, the prediction score for each class is defined as 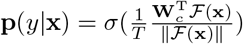, where *σ* indicates a softmax function. In our study, the sensitive and resistant prototypes are required to update their positions based on maximizing and minimizing entropy, we thus refer to the weight vectors W_*C*_ as the class prototypes.

We firstly train ℱ and 𝒞 using D_*s*_ and D_*t*_ so that the labeled examples can correctly classified. The standard cross-entropy is used to evaluate the classification loss:

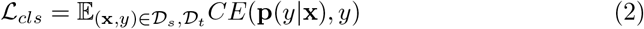

### Adversarial training

The adversarial training [56] introduces subtle perturbations within the vicinity of the input data. The perturbations are oriented towards the direction of loss gradient ascent and are typically generated under *l*_2_ norm constraints. This training strategy demands that the model not only minimizes the empirical risk but also minimizes the adversarial loss, making the model less sensitive to slight changes in the input. Formally, the adversarial loss is defined as below:

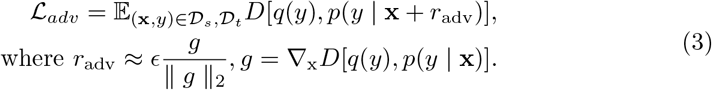

Note that *D*[·,·] represents the Kullback-Leibler divergence that measures the difference between two distributions, *p*(*y* | **x**) denotes the probability distribution of the predicted label *y* given input **x**, *q*(*y*) is the true distribution of label which is generally approximated by one-hot vector of *y, g* is the gradient that can be effectively computed using backpropagation, and *ϵ* is the magnitude of the adversarial perturbation. This perturbation strives to maximize the divergence between actual and predicted label distributions by choosing the direction of gradient ascent.

### Semi-supervised domain adaptation

The minimax entropy method [31] is adopted for semi-supervised domain adaptation, which maximizes the entropy of unlabeled target samples to optimize 𝒞, and minimizes the entropy with respect to ℱ to aggregate features. For the unlabeled samples, we assume the existence of a domain-invariant prototype in both the sensitive and resistant categories, representing the central points for two domains. Initially, ℱ and 𝒞 are trained using the labeled source samples and a small number of labeled target samples, and then the initial drug-sensitive and drug-resistant prototypes are evaluated. We utilize the unlabeled target samples to train 𝒞, maximizing the entropy between the prototypes and the features of unlabeled target samples. This allows us to update the prototype, and bring them closer to the unlabeled target samples. Next, to extract discriminative feature, we train ℱ to reduce the entropy between the prototypes and the unlabeled target features, thereby the features are clustered around one of the two prototypes in the latent space. The entropy is defined as below:

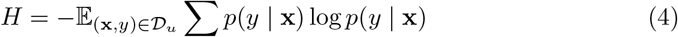

Therefore, the minimax entropy stands as an adversarial training process, which maximizes the entropy to find the prototypes that can best represent the unlabeled target features and then strives to minimize entropy to cluster the unlabeled features based on these prototypes. In this study we utilize the gradient reversal layer to implement the adversarial learning. Denote by ℒ_*mme*_ the loss of the minimax entropy-based adversarial learning, we have

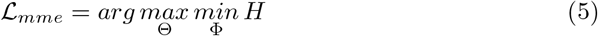

in which Θ and Φ are the parameters of classifier 𝒞 and feature extractor ℱ. As a result, the total loss function of our SSDA4Drug is defined as

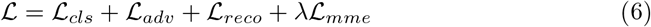

in which *λ* is a hyperparameter used for the trade-off between the minimax entropy loss and the classification task loss on the labeled samples. The detailed network architecture has been presented in Supplementary Table 3.

### Identification of key genes

We first consider the Integrated Gradients [51] to identify key genes. Integrated Gradients provides a quantitative assessment of the importance of each input feature in the decision-making, enhancing the interpretability of our model. In our practice, the integrated gradient (IG) scores were calculated using the “IntegratedGradients” function in the Python Captum library [57], taking as inputs the expression matrix, trained model and output labels. It output the IG matrices with the same shape as the input expression matrix, its rows represent genes and columns represent cells and elements are the corresponding IG values [22]. To identify genes with significantly higher IG values in sensitive (or resistant) cells, we performed differential expression analysis using the “rank genes groups” function. We considered the genes with logFC*>*1 and p-adj*<*0.05 as key genes.

We also leverage the SHAP values to uncover key genes involved in drug response. SHAP quantifies how much an input feature contributes to pushing the model’s prediction away from the base value, which is typically defined as the average output over the training dataset. We use the DeepExplainer from the SHAP [48] library to compute the SHAP values for each cell. To visualize the SHAP values, the “summary plot” function is used to plot the distribution of SHAP values for top 20 genes contributing significant to final prediction, i.e. resistance or sensitivity. The “beeswarm plot” function is used to plot the distribution of SHAP values within a specific group of cancer cells, such as drug-resistant cells.

### Pesudotime analysis

Monocle3 was used to analyze the evolutionary trajectory of different cell subgroups, using the gene-cell matrix in the UMI count scale extracted from the Seurat subset. The UMAP tool was used to reduce the dimensions. After dimensionality reduction and cell sorting, evolutionary trajectories were inferred using the default parameters of Monocle3.

## Declarations

### Funding

This work was supported by National Natural Science Foundation of China (No. 62072058, No. 62372229), Natural Science Foundation of Jiangsu Province (No. BK20231271).

### Conflict of interest

The authors declare no competing interests.

### Ethics approval and consent to participate Not applicable

### Data availability

GDSC is publicly available through the website (https://www.cancerrxgene.org/). Drug response annotation, including half maximal inhibitory concentration (IC50) and area under the dose-response curve (AUC), are available through the page https://www.cancerrxgene.org/downloads/bulk_download. Gene expression data (RMA-normalized basal expression profiles) for cell lines can be accessed on GDSC (https://www.cancerrxgene.org/gdsc1000/GDSC1000_WebResources/Home.html). The scRNA-seq data of MCF7 cell line under drug treatment at multiple time points was obtained from GEO with accession number GSE114462. The scRNA-seq data of PC9 cell line under drug treatment at multiple time points was obtained from GEO with accession number GSE149383.

The scRNA-seq data of PCa cell line was obtained from GEO with accession number GSE140440. The scRNA-seq data of patient-derived OSCC cells was obtained from GEO with accession number GSE117872. The scRNA-seq data of drug Etoposide and PLX4720 were obtained from GEO with accession number GSE149215 (GSM4494347: PC9 untreated; GSM4494354: PC9 cell treated with 25 µm Etoposide for 3 days) and GSE108383 (scRNA-seq for A375and 451Lu cell lines), respectively. The scRNA-seq of JHU006 and SCC47 cell line is available from Broad Institute’s single-cell portal (https://singlecell.broadinstitute.org/single_cell/study/SCP542/).

### Code availability

The source code and processed datasets of this study are available at GitHub: https://github.com/hliulab/SSDA4Drug.

### Author contribution

H.L. and K.H. conceptualized the idea. K.H. implemented the model. K.H. collected the data and conducted the experiments. K.H. and H.L. plotted figures. H.L. and K.H. prepared the manuscript. H.L. revised the manuscript. H.L. supervised the research.

## Notes

### Competing Interest Statement

The authors have declared no competing interest.

### Summary of Updates

1. More performance comparison experiments have been conducted. Figure 2 revised. 2. We also extended the Discussion section.

